# MEDUSA: Maintaining Entire DNA Duplexes for Utmost Sequencing Accuracy

**DOI:** 10.1101/2025.11.07.687202

**Authors:** Kan Xiong, Rachel Li, Ning Li, Ruolin Liu, Azeet Narayan, Justin Rhoades, Catherine Song, R. Coleman Lindsley, Heather A. Parsons, G. Mike Makrigiorgos, Viktor A. Adalsteinsson

## Abstract

Highly accurate DNA sequencing is essential for many applications. Duplex sequencing-based methods achieve unparalleled accuracy by requiring reads from both strands of the original DNA duplex to match. Yet, methods to prepare dsDNA for sequencing may resynthesize portions of each DNA duplex and cause base damage errors on one strand to become indistinguishable from true mutations on both strands. Here, we report MEDUSA (Maintaining Entire DNA Duplexes for Utmost Sequencing Accuracy) which minimizes dsDNA resynthesis to maximize duplex sequencing accuracy and yield. MEDUSA carefully repairs and blunts fragmented dsDNA, then employs apyrase to digest residual dNTPs, followed by restricted dA-tailing to prevent resynthesis. MEDUSA affords full genome coverage in a simplified protocol that is broadly compatible with dsDNA fragmentation and library preparation kits. We benchmarked MEDUSA on sheared genomic DNA and cell-free DNA and found a residual SNV frequency within a median 1.23-fold (range 0.92-1.88; p < 0.001) of what was expected if resynthesis was mostly blocked, but with full genome coverage and duplex yields within a median 1.02-fold (range 0.33 - 1.46; p = 0.258) of traditional methods that do not limit resynthesis. In all, MEDUSA could enable high breadth or depth of duplex sequencing while limiting false discovery.

**GRAPHICAL ABSTRACT:** 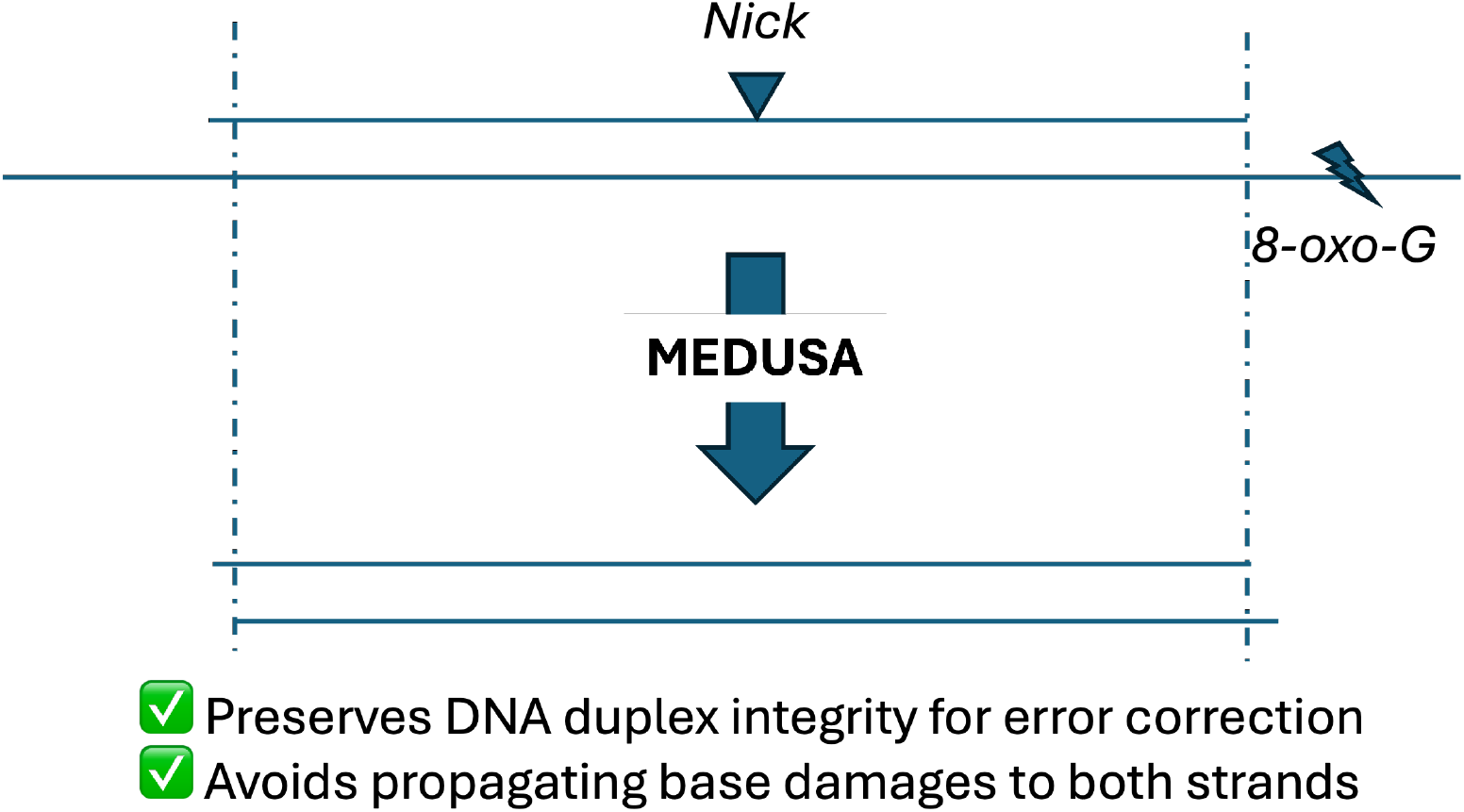

## INTRODUCTION

Highly accurate DNA sequencing is crucial for many applications in clinical care and research, including but not limited to cancer screening, precision medicine, prenatal testing, organ transplant monitoring, and studies of cancer evolution and somatic mosaicism (1–9). While the cost of DNA sequencing has declined dramatically over the past two decades, next generation sequencing (NGS) error rates have remained high (∼0.1%) (10, 11). This error rate makes it difficult to resolve true signals, particularly those which are present at low abundance. Higher accuracy sequencing is required for enhancing the performance of diagnostic tests and faithfully characterizing human and disease biology. One of the most accurate sequencing methods is duplex sequencing (12), which suppresses errors introduced during sample preparation and sequencing by requiring a true mutation to be detected on both strands of a duplex (13–15). However, the accuracy of duplex sequencing is highly dependent on DNA sample quality that can vary widely due to differences in sample collection and storage protocols (16, 17). During these processes, DNA base damages (e.g. uracil, thymine dimers, pyrimidine dimers, 8-oxoguanine [8-oxoG], depurination, depyrimidination.) and backbone damages (e.g. nicks, gaps or 5’ overhangs) can accumulate (18). These DNA lesions could introduce false mutations in duplex sequencing if the base damage errors are artificially propagated to both strands when the DNA duplex is prepared for ligation of NGS adapters (16, 19–21). Consequently, there is a strong need for a duplex sequencing method(s) that can afford high accuracy regardless of DNA sample quality while enabling high DNA duplex recovery and full genome coverage through a simplified workflow.

To prepare libraries for duplex sequencing, fragmented DNA such as cell-free DNA (cfDNA) or sheared genomic DNA (gDNA) are commonly subjected to End Repair and dA-Tailing (ER/AT) to facilitate ligation to NGS adapters. During ER/AT, DNA polymerases employed to fill in gaps and overhangs may resynthesize portions of each duplex, which has been known to occur near the ends of DNA fragments due to the presence of 5’ overhangs (13). When DNA lesion-induced mutations that are originally restricted to single strands are copied to both strands, these alterations manifest as duplex errors indistinguishable from bona fide mutations. The extent of strand resynthesis has been shown to be strongly associated with the residual SNV frequency (16). Lately, groups have developed ER/AT methods which minimize DNA resynthesis. For instance, NanoSeq uses a restriction enzyme to digest gDNA into short blunted fragments and then performs non-A dideoxynucleotide (ddBTP)-blocked ER/AT (22). As a result, NanoSeq could achieve an error rate of < 5e-9; however, this approach only applies for sequences containing the enzymatic restriction site, thus it can only analyze a small portion of the human genome and does not work well on fragmented dsDNA. To analyze fragmented dsDNA, mung bean nuclease can be used with NanoSeq to blunt dsDNA (23); however, this enzyme has very low blunting efficiency and thus limited DNA duplex recovery. We recently developed a method called Duplex-Repair that significantly limits strand resynthesis compared to commercial ER/AT kits and can detect residual SNV frequency as low as 9.2e-8 (16). Since Duplex-Repair does not require restriction digestion, it therefore does not limit the genomic regions that can be analyzed and works well on fragmented dsDNA. However, it does not fully eliminate strand resynthesis, rendering its residual SNV frequency higher than that of ddBTP-blocked ER/AT (24), and its lengthy protocol with bead cleanups limits dsDNA duplex recovery (16). In this work, we first developed an improved AT protocol which further restricts strand resynthesis (Duplex-Repair V2). We then developed MEDUSA, a streamlined ER/AT method for genome-wide, highly accurate duplex sequencing with high duplex recovery and similar hands-on time as traditional ER/AT methods.

## MATERIAL AND METHODS

### Clinical specimens

All patients provided written informed consent to allow the collection of blood and/or tumor tissue and analysis of genetic data for research purposes. Healthy donor blood samples were ordered from Research Blood Components or Boston Biosciences. Patients with breast cancer were prospectively identified for enrollment into an IRB-approved tissue analysis and banking cohort (Dana-Farber Cancer Institute [DFCI] protocol identifier 06-169). Blood samples were collected from participants (IRB-approved protocol 15-161) for the study on pre-existing hematopoietic stem cell mutations and therapy-related myeloid neoplasms following autologous stem cell transplantation in non-Hodgkin lymphoma.

### Plasma and whole blood processing

cfDNA was extracted from healthy donor plasma using the QIAsymphony DSP Circulating DNA Kit and quantified using the Quant-iT PicoGreen assay on a Hamilton STAR-line liquid handling system. gDNA was extracted from whole blood or blood cells using the QIAsymphony DSP DNA mini kit and quantified using the Quant-iT PicoGreen assay on a Hamilton STAR-line liquid handling system. Extracted gDNA was then Covaris sheared to an average fragment size of 150 bp, or restriction digested by HpyCH4V (NEB, R0620S)) and AluI (NEB, R0137S)) followed by 0.8x AMPure XP reverse size-selection and then 1.8x AMPure XP cleanup to yield an average fragment size of 150 bp.

### Duplex-Repair v2

Duplex-Repair v2’s ER/AT consists of five steps. In step 1, DNA is treated with an enzyme cocktail consisting of EndoIV (Cat. No. M0304S), Fpg (Cat. No. M0240S), UDG (Cat. No. M0280S), T4 PDG (Cat. No. M0308S), EndoVIII (Cat. No. M0299S) and ExoVII (Cat. No. M0379S) (all from NEB; use 0.2 ul each) in 1× NEBuffer 2 in the presence of 0.05 ug/ul BSA (total reaction volume = 20 ul) at 37°C for 30 min. In step 2, T4 PNK (Cat. No. M0201S; NEB; use 0.25 ul), T4 DNA polymerase (Cat. No. M0203S; NEB; use 0.25 ul), ATP (final concentration = 0.8 mM), and dNTP mix (final concentration of each dNTP = 0.5 mM) are added into the step 1 reaction mix and incubated at 37°C for another 30 min. In step 3, HiFi Taq ligase (Cat. No. M0647S; NEB; use 0.5 ul) and 10× HiFi Taq ligase buffer (use 1.5 ul) are spiked into the step 2 reaction mix and incubated on a thermal cycler that heats from 35 °C to 65 °C over the course of 45 min. The resulting products are purified by performing 3X AMPure XP bead cleanup and eluted in 16.2 ul of 10 mM Tris buffer. In step 4, the purified products are treated with apyrase (Cat. No. M0398S; NEB; use 0.5 uL) at 30 °C for 20 min followed by heat inactivation at 65 °C for 20 min. In step 5, Klenow fragment (3′→ 5′ exo-) (Cat. No. M0212L; NEB; use 1 uL), Taq DNA polymerase (Cat. No. M0273S; NEB; use 0.2 uL), and dATP (final concentration = 0.2 mM) are spiked into the step 4 reaction mix (total reaction volume = 20 uL) and incubated at room temperature for 30 min followed by 65 °C for 30 min. To prepare Duplex-Repair v2 libraries for conventional duplex sequencing, T4 DNA ligase (Cat. No. M0202L; NEB; use 2.5 uL), 5′-deadenylase (Cat. No. M0331S; NEB; use 0.5 ul), PEG 8000 (final concentration = 10% w/v), and custom dual-index duplex UMI adapters (IDT) are added to the step 5 reaction mix (total reaction volume = 55 ul), which is then incubated at room temperature for 1 h followed by a 1.2× AMPure XP bead cleanup. The purified products are then amplified by PCR.

### MEDUSA

MEDUSA’s ER/AT consists of three steps. In step 1, DNA is treated with an enzyme cocktail consisting of the same enzymes as in Duplex-Repair v2’s step 1 (optional), T4 DNA polymerase (use 0.5 uL), T4 PNK (use 0.5 ul) and HiFi Taq ligase (use 0.5 ul) in an integrated buffer consisting of 2 uL of 10x NEBuffer 2, 0.4 ug BSA, 1.2 uL of 10 mM dNTP mix, 2 uL of 10 mM ATP and 1.5 uL of 10x HiFi Taq ligase buffer with a total reaction volume of 25.7 uL. The reaction mix is incubated at 37 °C for 15 min followed by gradually ramping the temperature from 37 °C to 65 °C over the course of 15 min, and then incubated at 65 °C for 15 min. In step 2, apyrase (use 0.5 uL) is spiked into the step 1 reaction mix, which is then incubated at 30 °C for 10 min followed by heat inactivation at 65 °C for 20 min. In step 3, Klenow fragment (3′ → 5′ exo-) (use 1.5 uL), Taq DNA polymerase (use 0.3 uL), and 0.6 uL of 10 mM dATP are spiked into step 2 reaction mix, which is then incubated at room temperature for 15 min followed by 65 °C for 15 min. To prepare MEDUSA libraries for conventional duplex sequencing, T4 DNA ligase (use 2000 units), 5′-deadenylase (use 1 ul), PEG 8000 (final concentration = 10% w/v), and custom dual-index duplex UMI adapters (IDT) are added to the step 3 reaction mix (total reaction volume = 60.1 ul) which is then incubated at room temperature for 1 h followed by a 1.8× AMPure XP bead cleanup. The purified products are then amplified by PCR.

### ddBTP-blocked ER/AT

Restriction digested DNA is dA-tailed in NEBuffer 4 (total reaction volume = 15 uL) with Klenow fragment (3′ → 5′ exo-) (use 0.15 uL) and dATP/ddBTP (ddTTP, ddCTP and ddGTP; final concentration of each = 1 mM) at 37 °C for 30 min.

### CODEC workflow

CODEC library preparation is performed as previously described (24). In brief, 1.8x AMPure XP cleanup is performed on ER/AT products, which are then ligated to the CODEC adapter quadruplex. Strand displacing extension is performed using phi29 DNA polymerase (NEB, M0269L) at 30 °C for 20 min, followed by 0.75× AMPure XP cleanup. The product is amplified using KAPA HiFi HotStart ReadyMix (Roche, 07958935001) and xGen Library Amplification Primer Mix (IDT, 1077677) following the KAPA Library Amplification Kit’s protocol with 2 min of extension. After the PCR, a 0.65×AMPure XP cleanup is performed twice. For cfDNA, the second cleanup follows the double-size selection protocol with 0.5× and 0.65× volume ratios. CODEC libraries are sequenced on the Illumina NovaSeq 6000 System (S4 flow cell, 2 × 169 cycles) and NovaSeq X (10B flow cell, 2 × 151 cycles).

### Hybrid capture duplex sequencing

WGS libraries were constructed using four library preparation methods: MEDUSA, Duplex-Repair v2, Duplex-Repair, and the KAPA HyperPrep Kit with custom dual-index duplex UMI adapters (IDT). The libraries were then quantified using the Quant-iT dsDNA assay kit. Conventional duplex sequencing was performed following the previously published method (13, 16). In brief, hybrid capture (HC) using patient-specific probe panels (IDT) is performed using the xGen Hybridization and Wash Kit with xGen Universal Blockers (IDT). For each HC reaction, libraries are pooled up to 8-plex with each sample’s library mass equivalent to 25 times the DNA mass into library construction. 4 pmol of probe panel (IDT) is applied to each pool. After the first round of HC, libraries are amplified by 16 cycles of PCR and then carried through a second round of HC using halved amounts of human Cot-1 DNA, xGen Universal Blockers, and probes. After the second round of HC, libraries are amplified by 8-10 cycles of PCR. The final captured products were sequenced with a target raw depth of 10,000x per site per 20 ng DNA mass into library construction.

### Conventional duplex sequencing data analysis

The returned FASTQs were first aligned, deduplicated, and recalibrated following the GATK Best Practices “Data pre-processing for variant discovery” workflow. Next, UMIs were extracted using fgbio and raw sequencing reads were converted into duplex consensus molecules. Residual SNV frequency and duplex recovery were then estimated as previously described (13).

### CODEC data analysis

CODEC data were analyzed using the standard CODECsuite pipeline from BCL files to variant calls, as previously described (24). The pipeline incorporates widely used tools, including Illumina’s bcl2fastq for base calling, fgbio for UMI processing, samtools for alignment file handling, and Picard for metrics collection. Both raw and molecular consensus BAM files were aligned using BWA-MEM to the hg38 reference without the decoy sequences available here (https://data.smaht.org/resources/genome-related/genome-reference-and-related-data). To accommodate the unique read structure of CODEC, the CODECsuite pipeline integrates specialized steps for demultiplexing, adapter trimming, and variant calling. The CODECsuite source code is publicly available on GitHub (https://github.com/broadinstitute/TAG-public), and the pipeline is maintained on Dockstore (https://dockstore.org/workflows/github.com/broadinstitute/TAG-public/SingleSampleCODEC:CODEC?tab=info).

## RESULTS

### Degrading residual dBTPs by apyrase treatment prior to dA-tailing (AT) could further limit strand resynthesis and improve duplex sequencing accuracy

As Duplex-Repair uses DNA polymerases with strong strand displacement activity for AT(16), we wondered whether any trace amount of dBTP carryover after Duplex-Repair’s ER (consisting of steps 1-3; Fig. S1) and the following 3x AMPure XP cleanup could result in strand resynthesis on dsDNA containing residual backbone damages during AT. To test this,we designed a 100 bp dsDNA oligo with a 30 bp 5’ overhang and labeled the two strands with different fluorophores such that we could quantify individual fragment lengths during AT using capillary electrophoresis. We subjected this oligo to Duplex-Repair’s AT and observed that the majority of its top strand was extended to ∼ 100 bp (middle vs. top blue traces in Fig. 1B), confirming that dBTP carryover could result in strand resynthesis during AT if there are residual backbone damages such as nicks or gaps that had not been repaired.

**Figure 1.**
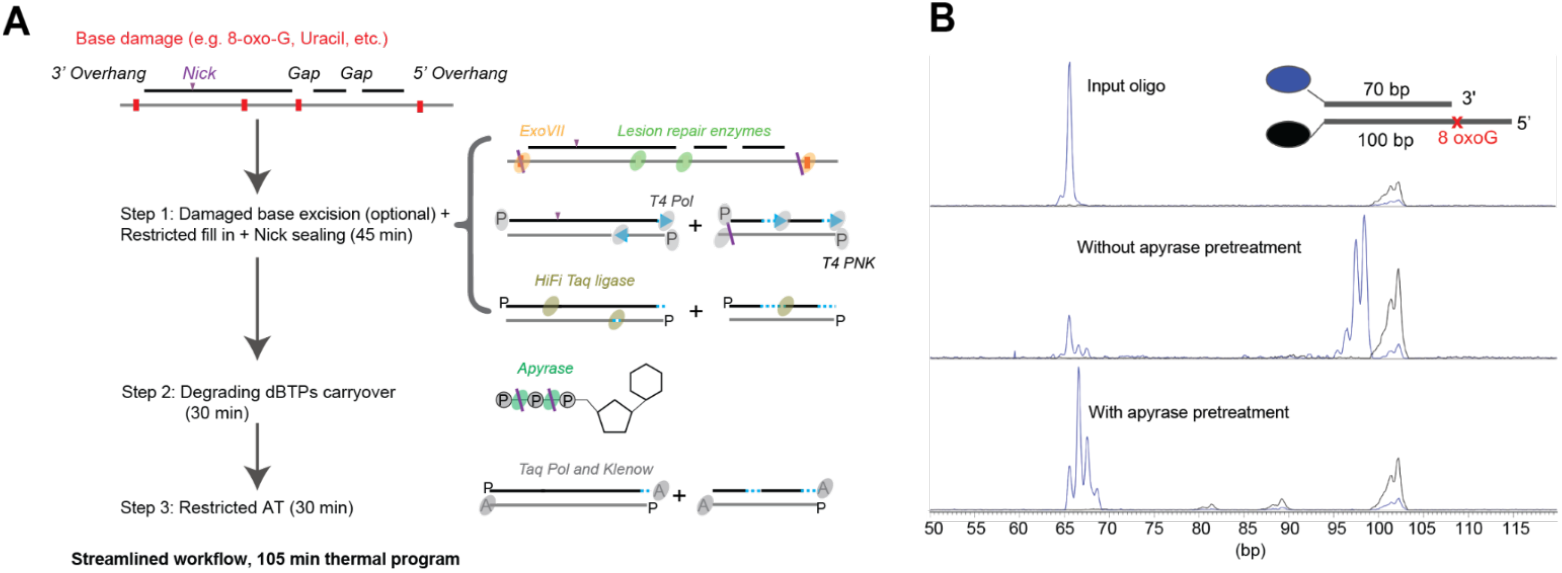
MEDUSA is a streamlined ER/AT method that could minimize strand resynthesis. A) Schematic of MEDUSA; B) Fragment analysis results of a synthetic duplex (annealed from oligos # 1 and 3 in Table S1) subjected to restricted dA-tailing with or without apyrase pretreatment.

To limit the potential for strand resynthesis during AT, we tested apyrase, an enzyme that hydrolyzes deoxynucleotide triphosphates to deoxynucleotide monophosphates (25), to degrade dBTP carryover prior to AT. We observed that pretreatment with 0.25 U of apyrase at 30 °C for 20 min sufficed to limit strand resynthesis during AT to <= 2 bp (bottom vs. top blue traces in Fig. 1B; Fig. S2). We further confirmed that apyrase treatment did not impact the efficiency of AT or downstream adapter ligation (Fig. S3). Based on these observations, we devised a new AT method that incorporated an additional apyrase treatment step (Steps 2 & 3 in Fig. 1A).

To assess whether this new AT method could further improve duplex sequencing accuracy, we treated three Covaris-sheared formalin fixed paraffin embedded (FFPE) tumor gDNA and three healthy donor (HD) cfDNA samples with Duplex-Repair, an ER/AT workflow combining Duplex-Repair’s ER and the new AT method (Duplex-Repair v2; Fig. S1), or the KAPA HyperPrep kit’s ER/AT (hereafter referred to as “standard ER/AT”), followed by hybrid capture duplex sequencing. For FFPE samples, Duplex-Repair v2 measured a median 2.61-fold (range 2.27-3.07) and 7.07-fold (range 7.01-10.92) lower residual SNV frequency than Duplex-Repair and standard ER/AT, respectively, with the most significant reduction in the C>T context (Fig. 2A & Fig. S4). Duplex-Repair v2 showed intermediate duplex recovery between Duplex-Repair and standard ER/AT (Fig. S5). For the three cfDNA samples, the reduction of residual SNV frequency by Duplex-Repair v2 over Duplex-Repair was limited (Fig. 2A). This result is consistent with previous observations that the impact of limiting strand resynthesis on duplex sequencing accuracy is dependent on DNA quality, with more damaged DNA such as FFPE gDNA standing to benefit more (16). Duplex-Repair v2 showed similar duplex recovery as Duplex-Repair (Fig. S6).

**Figure 2.**
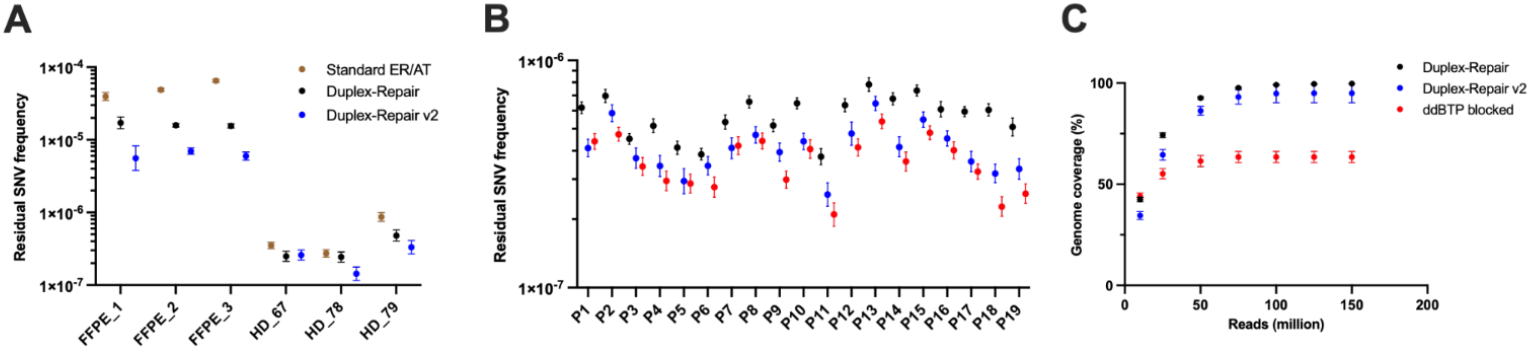
A new AT method could further improve duplex sequencing accuracy. A). Residual SNV frequencies of three FFPE tumor DNA and three cfDNA samples (two replicates per condition per sample) treated with standard ER/AT, Duplex-Repair, or Duplex-Repair v2, followed by hybrid capture duplex sequencing. B). Residual SNV frequencies of 19 samples treated with Duplex-Repair, Duplex-Repair v2, or ddBTP-blocked ER/AT, followed by CODEC whole genome duplex sequencing; C). CODEC genome coverages of 8 representative samples. Of note, Duplex-Repair v2 combines Duplex-Repair’s ER and a new AT method (Fig. S1).

### A new dA-tailing method could achieve similar residual SNV frequency as ddBTP- blocked ER/AT while allowing full genome coverage

We then assessed whether Duplex-Repair v2 could achieve similar residual SNV frequency as ddBTP-blocked ER/AT while affording full genome coverage and compatibility with fragmented DNA samples. To do so, we first treated 19 fragmented blood cell-derived gDNA samples with Duplex-Repair, Duplex-Repair v2, or ddBTP-blocked ER/AT, and then prepared Concatenating Original Duplex for Error Correction (CODEC) libraries for whole genome sequencing (WGS) (24). Of note, the DNA input into Duplex-Repair’s and Duplex-Repair v2’s ER/AT was Covaris sheared, which could introduce additional DNA damage (26), while the input into ddBTP-blocked ER/AT was restriction-digested. Restriction enzyme-based fragmentation kits that afford full genome coverage exist and could be used with Duplex-Repair, but ddBTP-blocked ER/AT is only compatible with enzymes that yield blunt-ended DNA (22). Following ER/AT, we applied CODEC, a library construction method that links both strands of each original DNA duplex such that they can be sequenced using one NGS read pair. As such, CODEC enables WGS with duplex sequencing accuracy using up to 100-fold fewer reads than conventional duplex sequencing (24).

We observed that across all samples, Duplex-Repair v2 and ddBTP-blocked ER/AT measured median 1.40-fold (range 1.12-1.91) and 1.53-fold (range 1.27-2.67) lower residual SNV frequency (median: 4.11e-7, range: 2.57e-7 – 6.46e-7; median: 3.59e-7, range: 2.1e-7 – 5.39e-7) than Duplex-Repair (median: 6.06e-7; range: 3.77e-7 – 7.83e-7), respectively (Fig. 2B). Duplex-Repair v2 measured slightly higher residual SNV frequency (median 1.15-fold higher, range 0.94-1.40-fold; p < 0.001, weighted paired t-test) than ddBTP-blocked ER/AT. We further examined context specific residual SNV frequencies and found that Duplex-Repair v2 and ddBTP-blocked ER/AT showed the largest residual SNV frequency reduction in the C>G and T>A contexts compared to Duplex-Repair (Fig. S7). In the C>G context, Duplex-Repair v2 and ddBTP-blocked ER/AT measured median 2.24-fold (range 1.70-5.11) and 5.87-fold (range 2.26-21.3) lower residual SNV frequency than Duplex-Repair, respectively. In the T>A context, Duplex-Repair v2 and ddBTP-blocked ER/AT measured 2.24-fold (range 1.19-5.29) and 2.35-fold (range 1.5-5.70) lower residual SNV frequency than Duplex-Repair, respectively. Duplex-Repair v2 measured higher residual SNV frequency in the C>G context (median 2.15-fold higher, range 1.01-7.33) than ddBTP-blocked ER/AT, possibly due to base damage errors introduced during Covaris shearing together with trace amounts of resynthesis that resulted in false C>G mutations on both strands. We next compared the human genome coverage of the three ER/AT methods by in silico downsampling of the CODEC data, and observed that ddBTP-blocked ER/AT could only cover up to 63.5% of the mappable human genome as expected from the combination of restriction enzymes used. Meanwhile, both Duplex-Repair and Duplex-Repair v2 could cover up to ∼ 100% of the mappable human genome (Fig. 2C). These results suggest that the new AT method could achieve residual SNV frequencies comparable to those of ddBTP-blocked ER/AT while allowing full genome coverage.

### MEDUSA is a streamlined and highly accurate ER/AT method

While Duplex-Repair v2 can achieve higher duplex sequencing accuracy than standard ER/AT, its workflow is longer and more complex than commercially-available protocols (Fig. S1). We therefore sought to streamline Duplex-Repair v2’s workflow. We hypothesized that the 3x AMPure XP cleanup included in Duplex-Repair (and v2) to remove excess dNTPs and limit strand resynthesis during AT could be omitted, given that Duplex-Repair v2 uses apyrase to degrade dNTPs. To test this, we subjected a synthetic oligo to Duplex-Repair v2’s ER/AT with or without the 3x AMPure XP cleanup, and observed that omitting the cleanup did not significantly impact AT and in fact improved DNA recovery (Fig. S8). We next assessed whether the incubation time for each step of Duplex-Repair v2’s ER/AT could be shortened. We found that halving the incubation time did not significantly impact the efficiency of ER/AT or downstream adapter ligation (Figs. S9 & S10). Furthermore, we evaluated whether Duplex-Repair v2’s three individual ER steps could be integrated into a single step. We first validated that several enzymes in the base excision repair (BER) step, in particular, formamidopyrimidine [fapy]-DNA glycosylase (Fpg), uracil-DNA glycosylase (UDG), and exonuclease VII (ExoVII), were still active in the new integrated ER buffer (Figs. S11-S13). Of note, ExoVII treatment left a ∼ 4 bp longer 5’ overhang in the integrated ER buffer than in the BER buffer (Fig. S13), which could elevate strand resynthesis during restricted fill-in by T4 DNA polymerase. Overall, we observed that integrating Duplex-Repair v2’s ER steps into a single step did not significantly impact the ER efficiency (Fig. S14). Based on these results, we devised a streamlined ER/AT method named Maintaining Entire DNA Duplexes for Utmost Sequencing Accuracy (MEDUSA) that consists of three steps: 1) BER (optional), restricted fill-in, and nick sealing, 2) degrading dBTP carryover, and 3) restricted dA-tailing (Fig. 1A). BER is deemed optional for higher quality DNA such as fresh frozen DNA and fresh cfDNA but recommended for lower quality DNA such as FFPE DNA and archival cfDNA. MEDUSA’s ER/AT could be completed in ∼ 2 hr, in contrast with more than 4 hr required for Duplex-Repair v2 (Fig. S1).

To assess the accuracy of MEDUSA, we selected 11 out of the 19 blood cell-derived gDNA samples analyzed in Fig. 2B, which were previously Covaris-sheared. We treated these DNA samples with MEDUSA or standard ER/AT and then prepared CODEC libraries for WGS. Across all samples, MEDUSA and ddBTP-blocked ER/AT measured median 2.84-fold (range 2.51 - 4.36) and 3.85-fold (range: 2.98 - 7.47) lower residual SNV frequency (median: 4.5e-7, range: 3.53e-7 – 8.77e-7; median: 3.96e-7, range: 2.25e-7 – 5.13e-7) than standard ER/AT (median: 6.06e-7; range: 3.77e-7 – 7.83e-7), respectively (Fig. 3A). MEDUSA measured slightly higher residual SNV frequency (median 1.23-fold higher, range 0.92-1.88) than ddBTP-blocked ER/AT. We further examined context specific residual SNV frequencies and found that MEDUSA and ddBTP-blocked ER/AT showed the largest reduction in residual SNV frequency in the C>A and C>G contexts compared to standard ER/AT (Fig. S15). In the C>A context, MEDUSA and ddBTP-blocked ER/AT measured a median 13.36-fold (range 9.27-29.74) and 19.56-fold (range 3.96 - 38.11) lower residual SNV frequency than standard ER/AT, respectively. In the C>G context, MEDUSA and ddBTP-blocked ER/AT measured a median 2.71-fold (range 1.28-4.14) and 8.88-fold (range 2.42 - 23.79) lower residual SNV frequency than standard ER/AT, respectively. Consistent with the observation above, MEDUSA measured higher background SNV frequency in the C>G context (median 3.07-fold, range 1.27-7.62) than ddBTP-blocked ER/AT.

**Figure 3.**
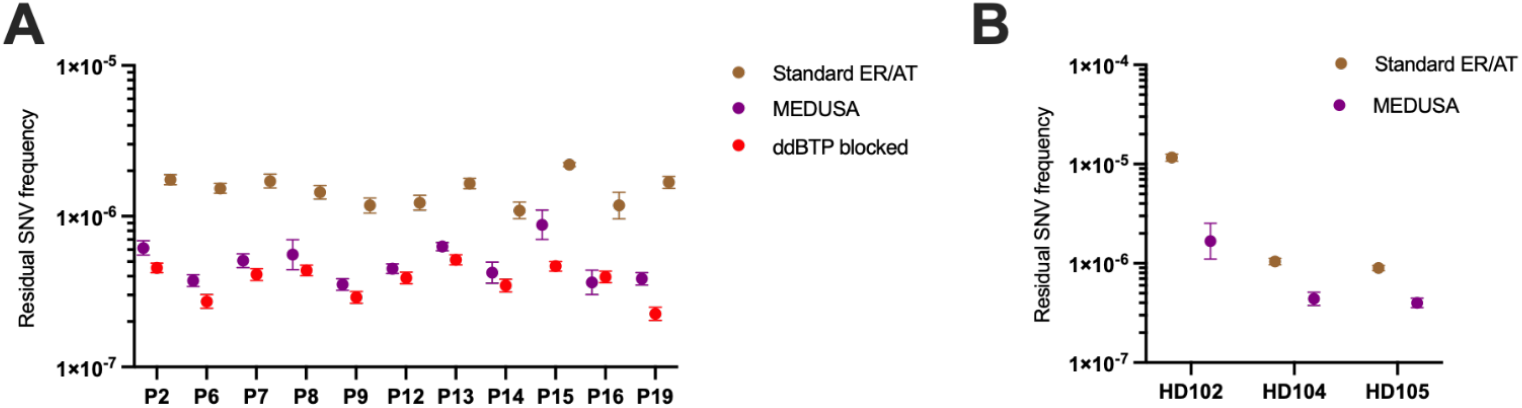
MEDUSA is a streamlined and highly accurate ER/AT method. A). Residual SNV frequencies of 11 samples treated with standard ER/AT, MEDUSA, or ddBTP-blocked ER/AT, followed by CODEC whole genome duplex sequencing. Of note: the residual SNV frequencies measured by ddBTP-blocked ER/AT from Fig. 2B were plotted here to facilitate comparison; B). Residual SNV frequencies of 3 HD cfDNA samples treated with standard ER/AT or MEDUSA, followed by hybrid capture duplex sequencing.

We further compared the background SNV frequencies of MEDUSA vs. Duplex-Repair v2 and found MEDUSA measured similar residual SNV frequency (median 1.05-fold, range 0.8-1.60; p = 0.177, weighted paired t-test) as Duplex-Repair v2 (Fig. S16). As expected, MEDUSA could cover up to ∼ 100% of the mappable human genome, while ddBTP-blocked ER/AT with the combination of restriction enzymes chosen could only cover up to 66.7% of the human genome (Fig. S17). We also treated three healthy donor cfDNA samples with MEDUSA or standard ER/AT and then performed hybrid capture duplex sequencing. We observed that MEDUSA measured a median 2.34-fold (range 2.24-6.94) lower residual SNV frequency than standard ER/AT (Fig. 3B). MEDUSA could recover similar duplex depths as standard ER/AT (median 1.02 fold, range 0.33 - 1.46; p = 0.258, Wilcoxon test), though its variability appeared higher than that of standard ER/AT (Fig. S18). These results indicate that MEDUSA enables highly accurate sequencing with full genome coverage and high molecular recovery.

## DISCUSSION

We have developed a highly accurate and streamlined ER/AT method that limits DNA resynthesis and propagation of mutations arising from base damage errors to both strands of each DNA duplex, while affording full genome coverage, compatibility with fragmented DNA, and high molecular recovery. To achieve this, we first strove to attain similar accuracy as NanoSeq, or ddBTP-blocked ER/AT. We found that trace amounts of dBTP carryover could result in strand resynthesis during AT if there were unresolved backbone damages. Based on this insight, we devised a new AT method that incorporates an apyrase treatment step to degrade dBTP carryover and thus limit strand resynthesis. We showed that this new AT method could achieve similar duplex sequencing accuracy as ddBTP-blocked ER/AT while allowing full genome coverage. Ultimately, we incorporated these improvements into a streamlined ER/AT workflow named MEDUSA, and showed that MEDUSA could achieve significantly higher duplex sequencing accuracy than standard ER/AT while maintaining full genome coverage and comparable duplex recovery efficiency.

The discovery that dBTP carryover could result in significant strand resynthesis during AT came as a surprise, as we expected the 3x AMPure cleanup between Duplex-Repair’s ER and AT to remove most, if not all, of the dBTPs used for ER. This finding may have implications for other workflows requiring dNTP removal. Moreover, though MEDUSA is more streamlined than Duplex-Repair v2, it still requires additional steps as compared to standard ER/AT. In particular, the reaction tube must be opened twice in order to add reagents, but MEDUSA otherwise has a similar hands-on time (∼20 min vs ∼10 min) and only a slightly longer overall time (∼120 min vs ∼70 min) than standard ER/AT. Future work will involve further streamlining MEDUSA into a one-pot reaction to enable more consistent duplex recovery. Additionally, pre-made stocks of MEDUSA’s enzyme and/or buffer mixtures could be used to minimize variability during pipetting and mixing.

MEDUSA could expand the applicability and feasibility of duplex sequencing approaches that demand high accuracy, high breadth or depth of genomic coverage, and compatibility with a wide range of sample types. MEDUSA can be used with different dsDNA fragmentation approaches, such as Covaris or enzymatic shearing, or with inherently fragmented samples such as cfDNA. For DNA samples that have been collected and stored under various conditions, including repeated freeze and thaw cycles, MEDUSA can limit errors resulting from base and backbone damages and thereby reduces the impact of pre-analytic factors. Given its high dsDNA recovery, MEDUSA is suitable for targeted deep sequencing of limited specimens such as liquid biopsies or fine needle aspiration biopsies. Furthermore, MEDUSA can be integrated with novel NGS library preparation methods such as CODEC (24) and Methyl-CODEC (27) for high breadth or depth sequencing. Although our efforts in this study focused on the coupling of MEDUSA with Illumina sequencing, it could in principle be used with other next- or third-generation sequencing technologies that involve dsDNA adapter ligation.

In summary, we developed MEDUSA, a highly accurate ER/AT method that minimizes dsDNA resynthesis, maximizes dsDNA recovery, and affords full genome coverage in a streamlined workflow. We envision that MEDUSA could enable high accuracy, breadth, and depth of duplex sequencing for a broad range of samples and applications.

## ACKNOWLEDGEMENTS

The authors would like to acknowledge the patients and their families for their contributions to this study.

## AUTHOR CONTRIBUTIONS

K. Xiong: Conceptualization, Investigation, Methodology, Visualization, Writing—original draft. R. Li: Investigation, Writing—review & editing. N. Li, J. Rhoades and C. Song: Formal analysis. R. Liu, A. Narayan, R. C. Lindsley and H. A. Parsons: Resources. G. M. Makrigiorgos: Writing—review & editing, funding acquisition. V. A. Adalsteinsson: Supervision, Conceptualization, Writing—original draft, funding acquisition.

## SUPPLEMENTARY DATA

Supplementary Data are available at NAR online.

## DATA AVAILABILITY

All sequencing data will be available in dbGaP.

