## Supplementary Data for "MEDUSA: Maintaining Entire DNA Duplexes for Utmost Sequencing Accuracy"

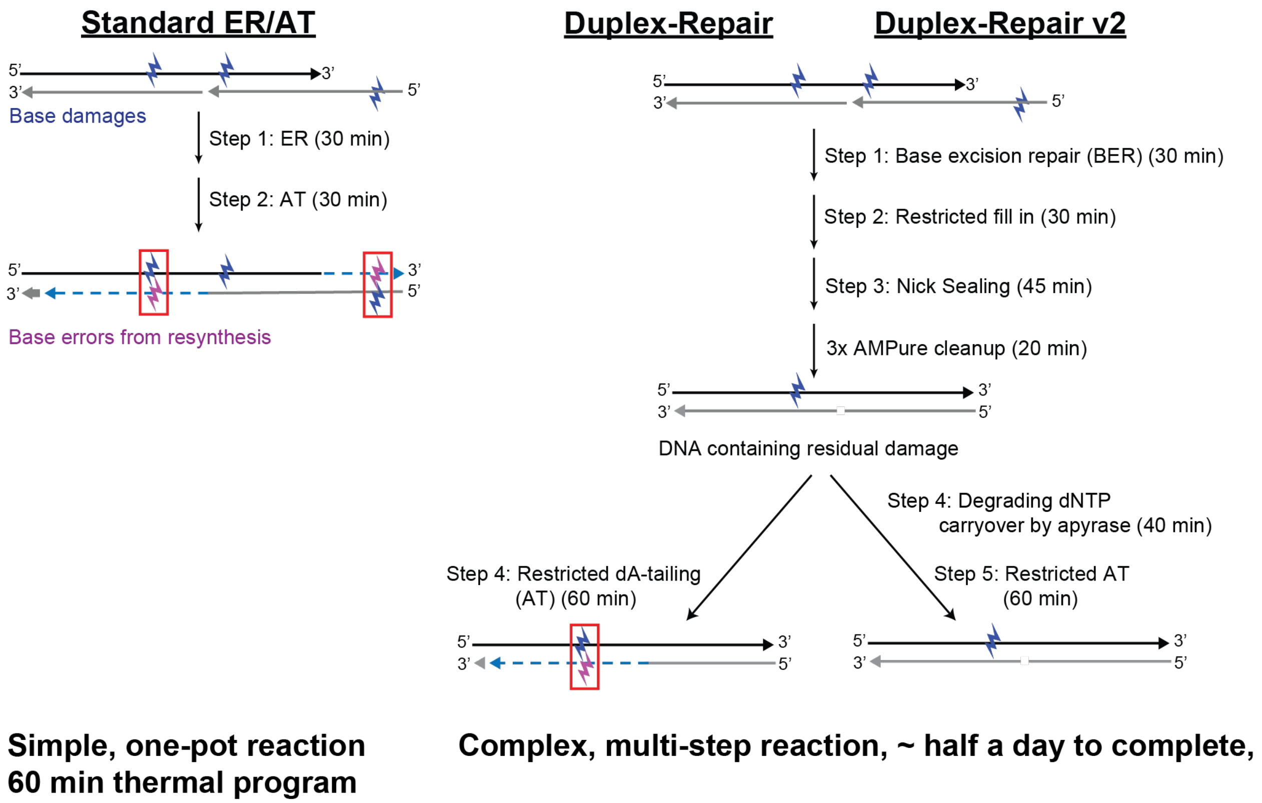


**Figure S1**: Schematics of standard ER/AT, Duplex-Repair and Duplex-Repair v2.

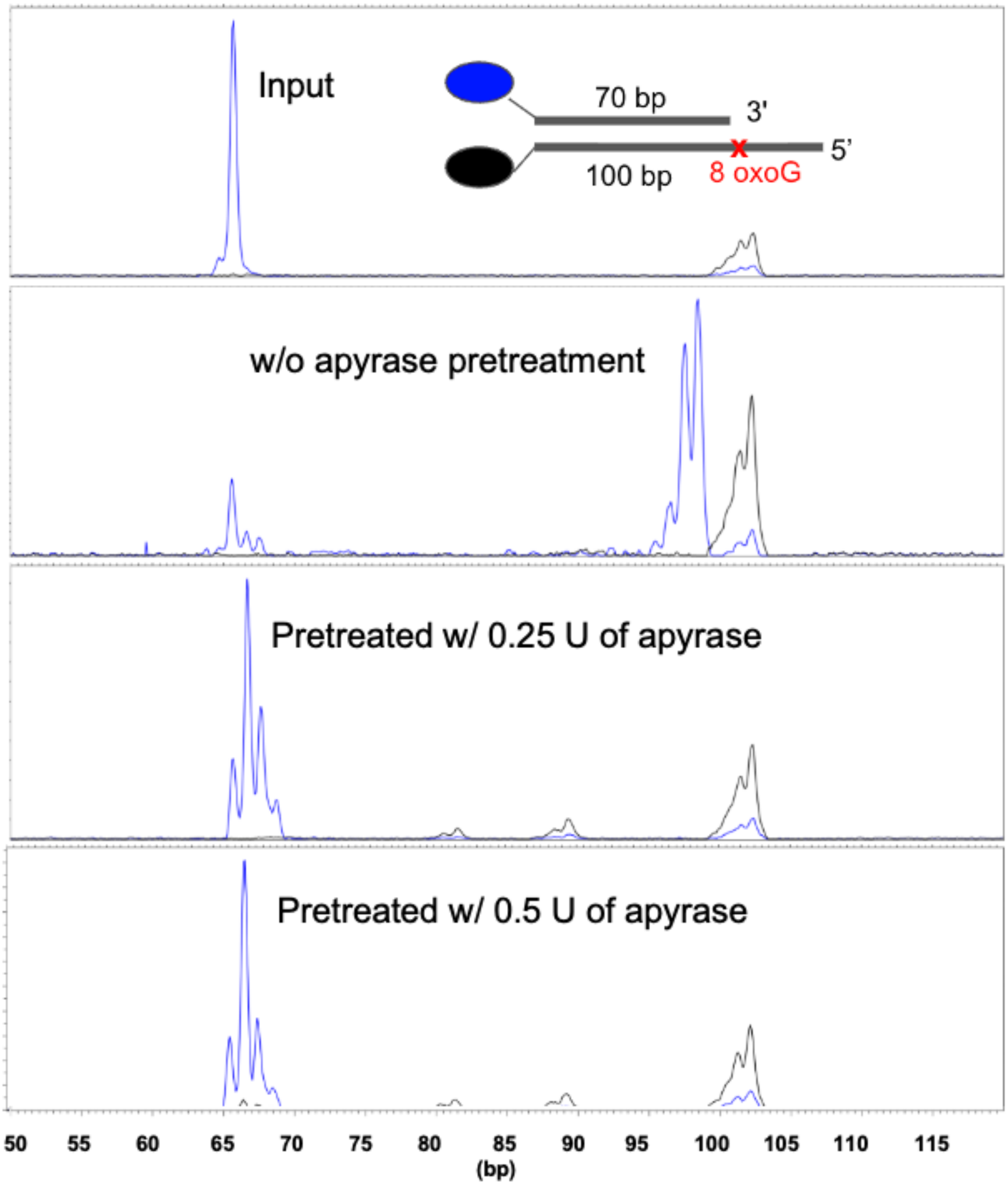


**Figure S2**: Fragment analysis results of a synthetic duplex subjected to restricted dA-tailing with or without apyrase pretreatment. Pretreatment with 0.25 or 0.5 U of apyrase for 20 min sufficed to limit strand resynthesis during restricted dA-tailing.

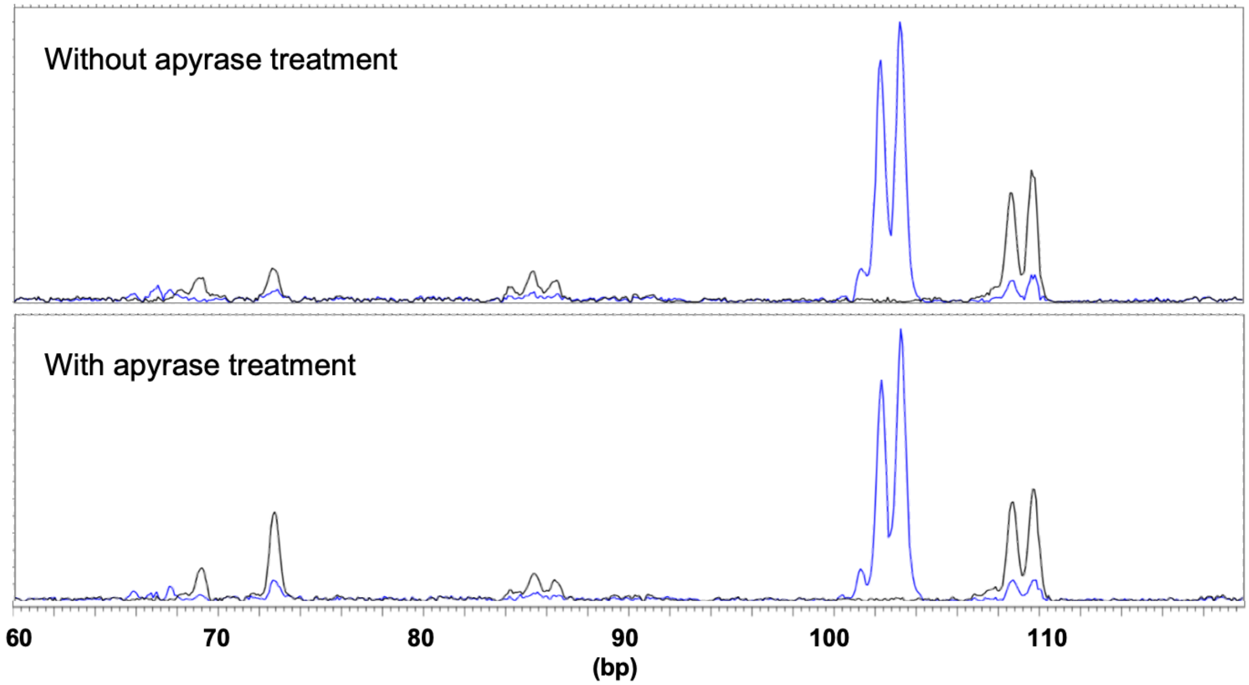


**Figure S3**: Fragment analysis results of a synthetic duplex subjected to Duplex-Repair's ER/AT with or without apyrase treatment, followed by adapter ligation. Apyrase treatment did not impact downstream dA-tailing and adapter ligation. The same synthetic duplex as in Fig. 1B and Fig. S1 was used. Of note, the base excision repair step was skipped.

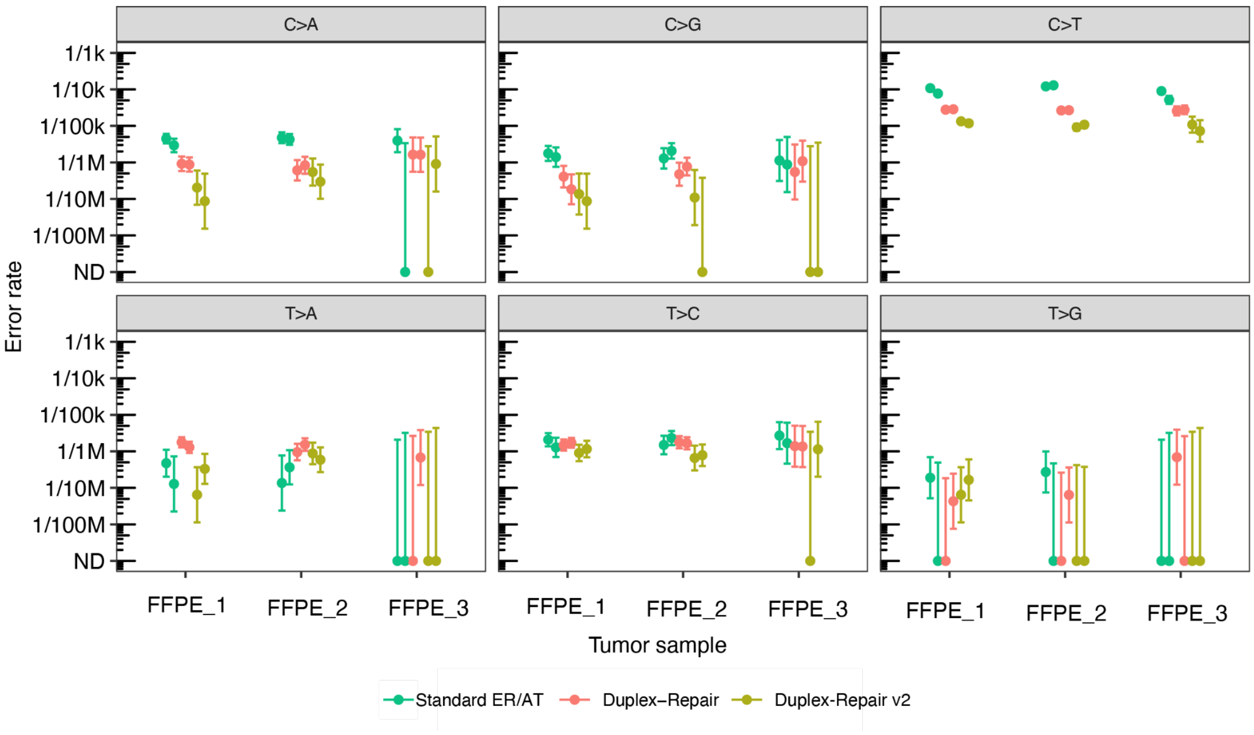


**Figure S4**: Context specific duplex error rates of three FFPE tumor biopsies (two replicates per condition per sample) treated with standard ER/AT, Duplex-Repair or Duplex-Repair v2, followed by hybrid capture duplex sequencing.

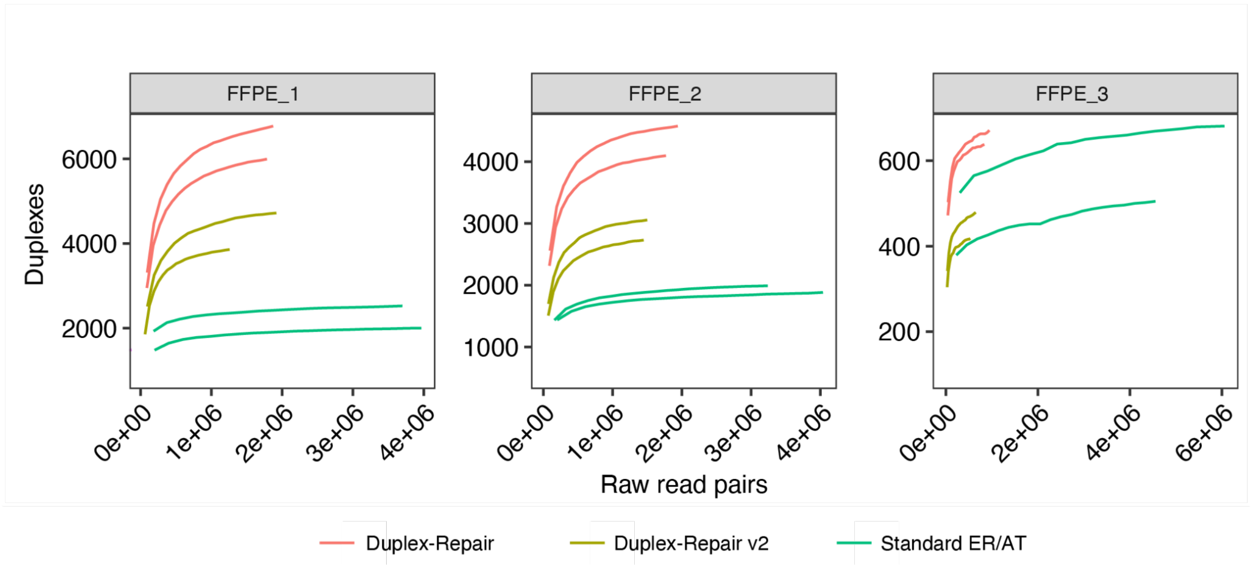


**Figure S5**: Total duplexes covered of three FFPE tumor biopsies (two replicates per condition per sample) treated with standard ER/AT, Duplex-Repair or Duplex-Repair v2, followed by hybrid capture duplex sequencing.

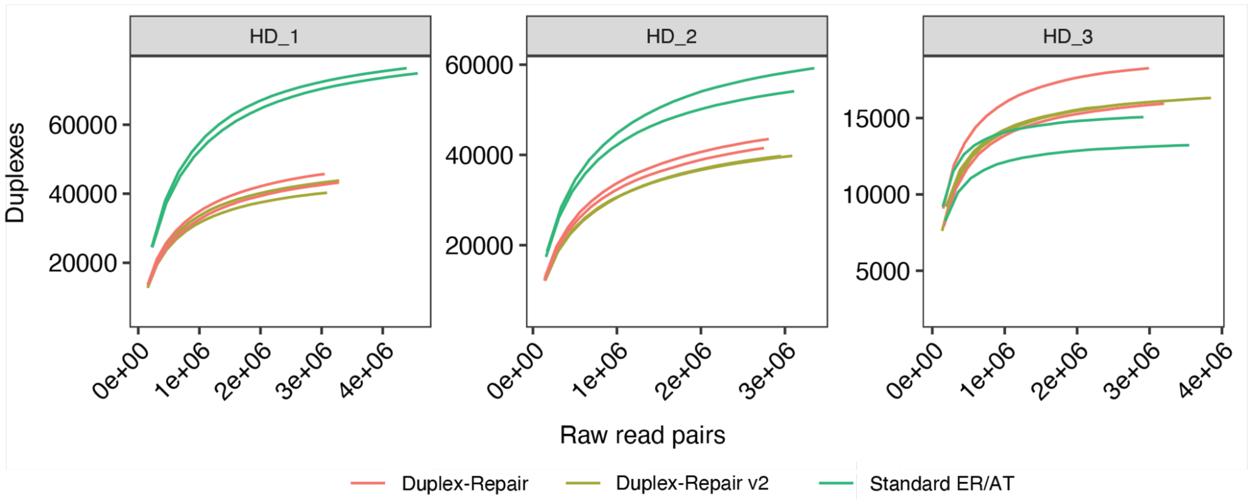


**Figure S6:** Total duplexes recovered of three healthy donor cfDNA (two replicates per condition per sample) treated with standard ER/AT, Duplex-Repair or Duplex-Repair v2, followed by hybrid capture duplex sequencing.

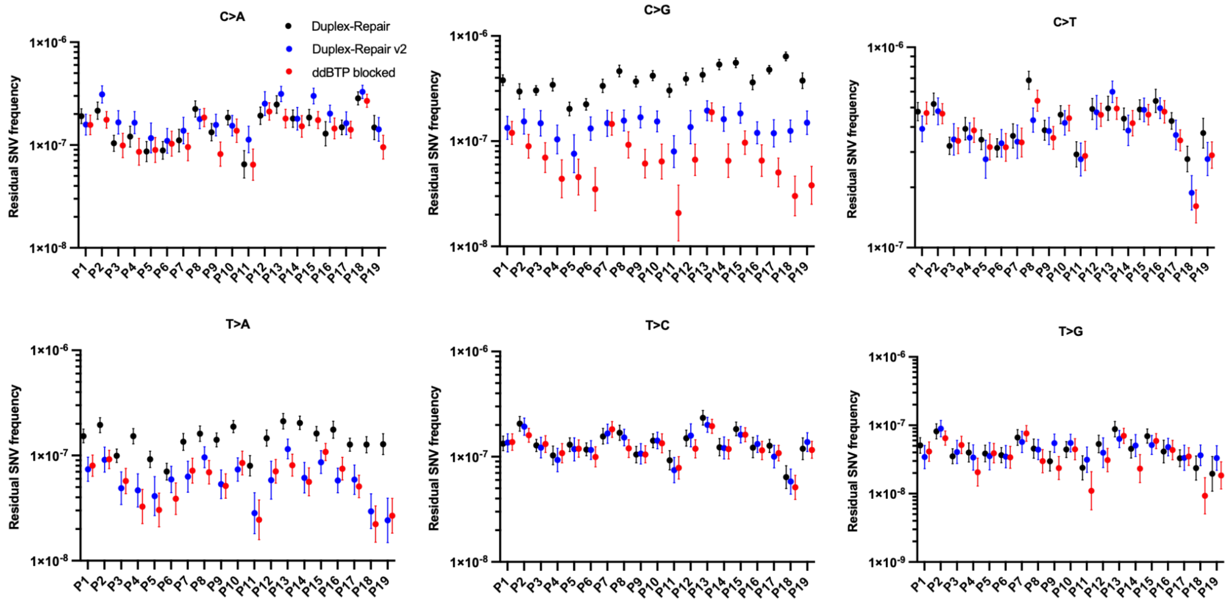


**Figure S7:** Context specific residual SNV frequencies of 19 samples treated with Duplex-Repair, Duplex-Repair v2 or ddBTP-blocked ER/AT, followed by CODEC whole genome duplex sequencing.

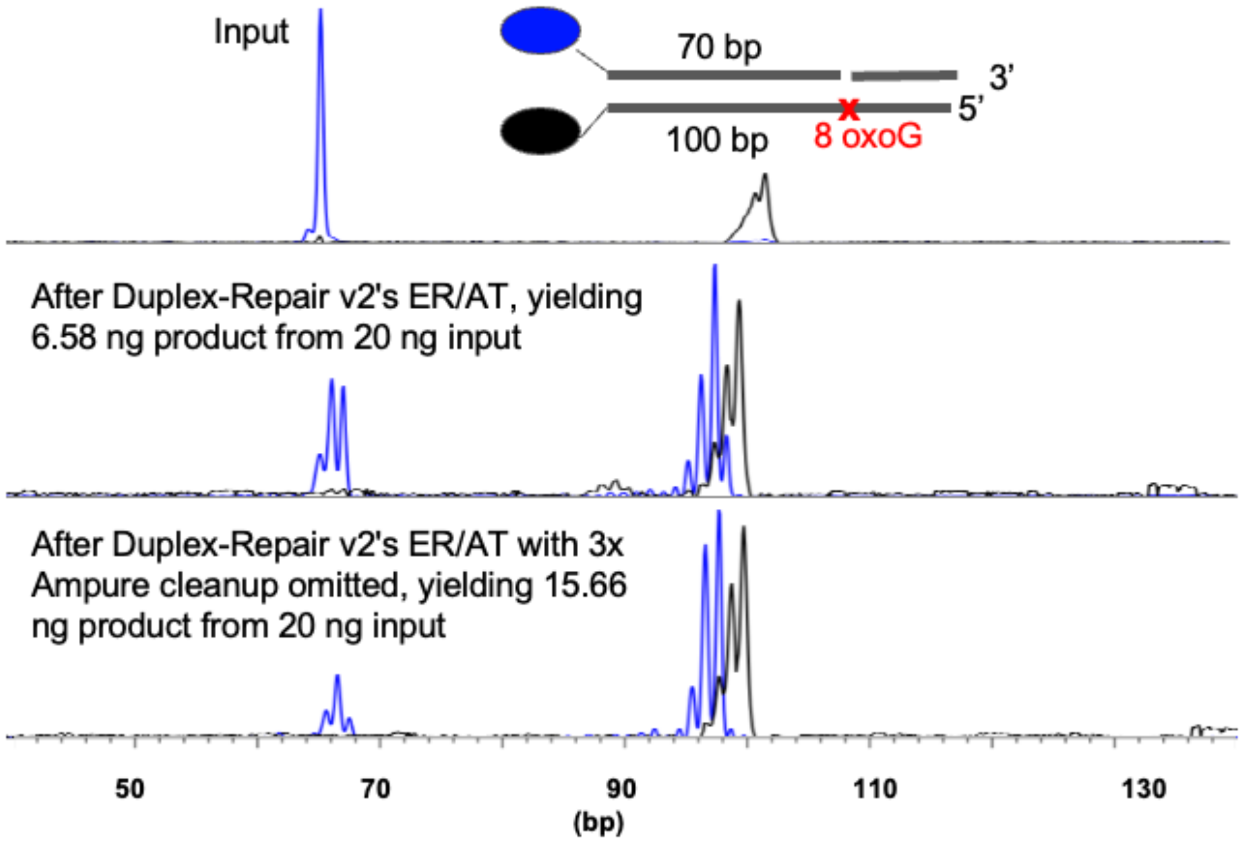


**Figure S8:** Fragment analysis results of a synthetic duplex (annealed from oligos # 1, 2 and 3 in Table S1) subjected to Duplex-Repair v2's ER/AT. Omitting the 3x AMPure cleanup in Duplex-Repair v2's workflow did not significantly impact the downstream dA-tailing. Of note, the BER step was skipped.

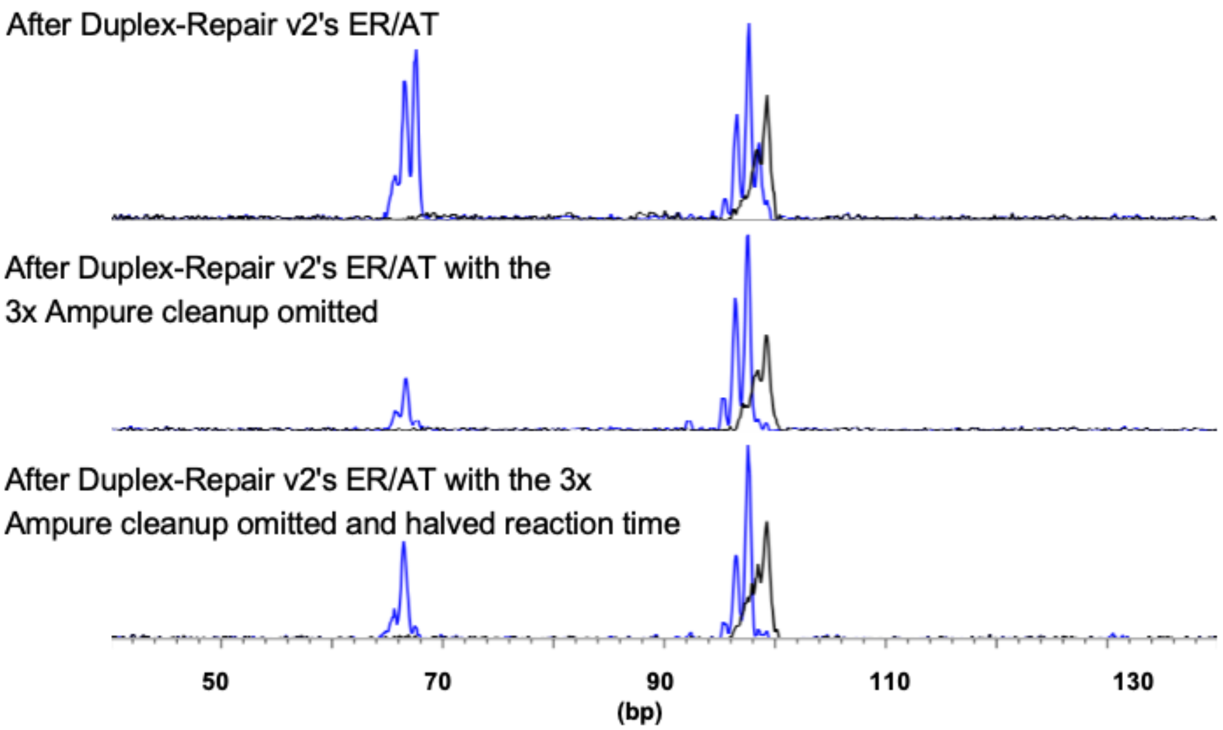


**Figure S9:** Fragment analysis results of a synthetic duplex subjected to Duplex-Repair v2's ER/AT. Halving the reaction time did not significantly impact Duplex-Repair v2's ER/AT efficiency. The same synthetic duplex as in Fig. S8 was used. The DNA input was 10 ng, and the yields of dA-tailed product from the three conditions (two replicates each) below were 2.41 ± 0.09, 3.51 ± 0.21 and 3.24 ± 0.15 ng, respectively. Of note, the BER step was skipped.

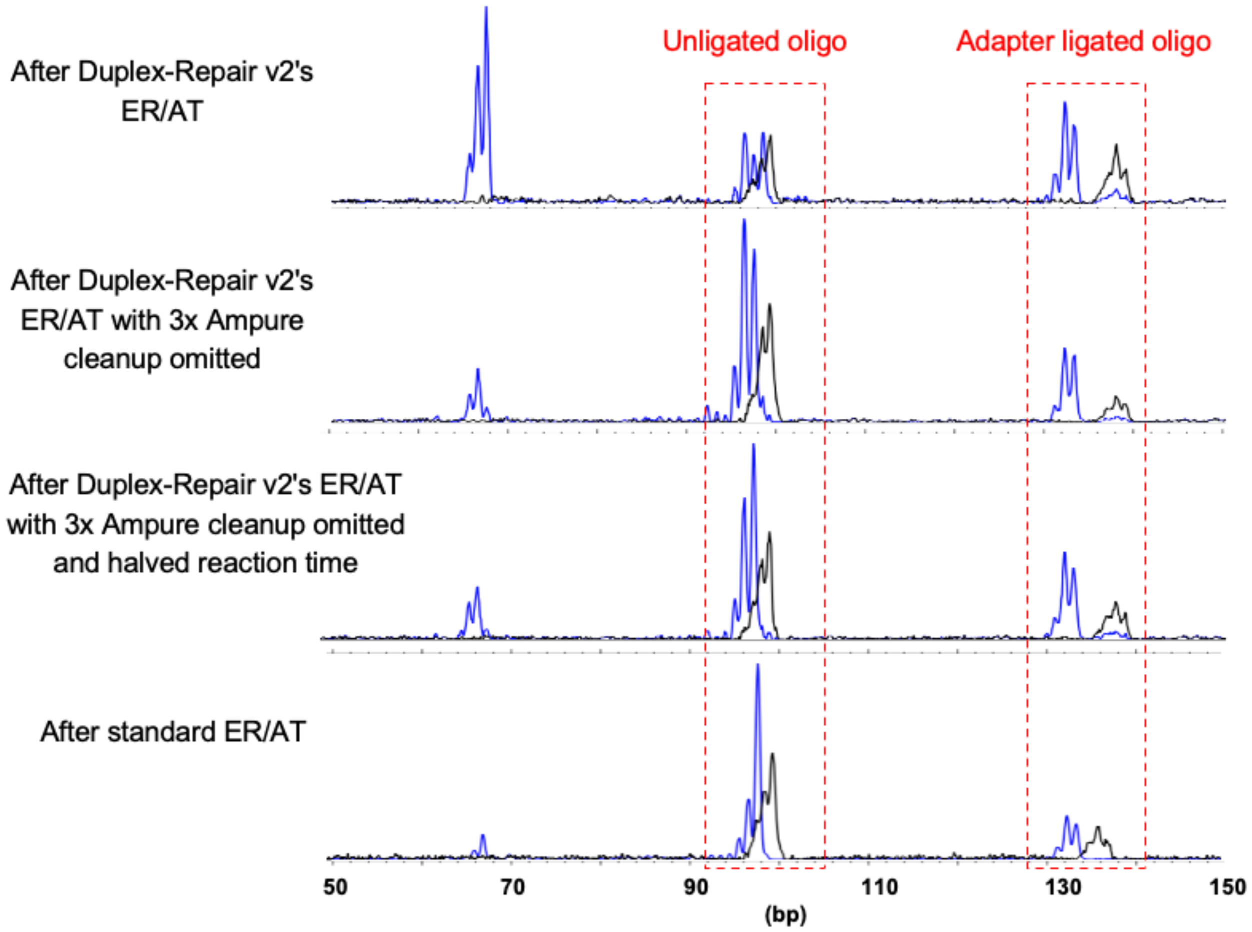


**Figure S10**: Fragment analysis results of a synthetic duplex subjected to Duplex-Repair v2's ER/AT followed by adapter ligation. The same synthetic duplex as in Fig. 1B and Fig. S2 was used. Of note, the base excision repair step was skipped.

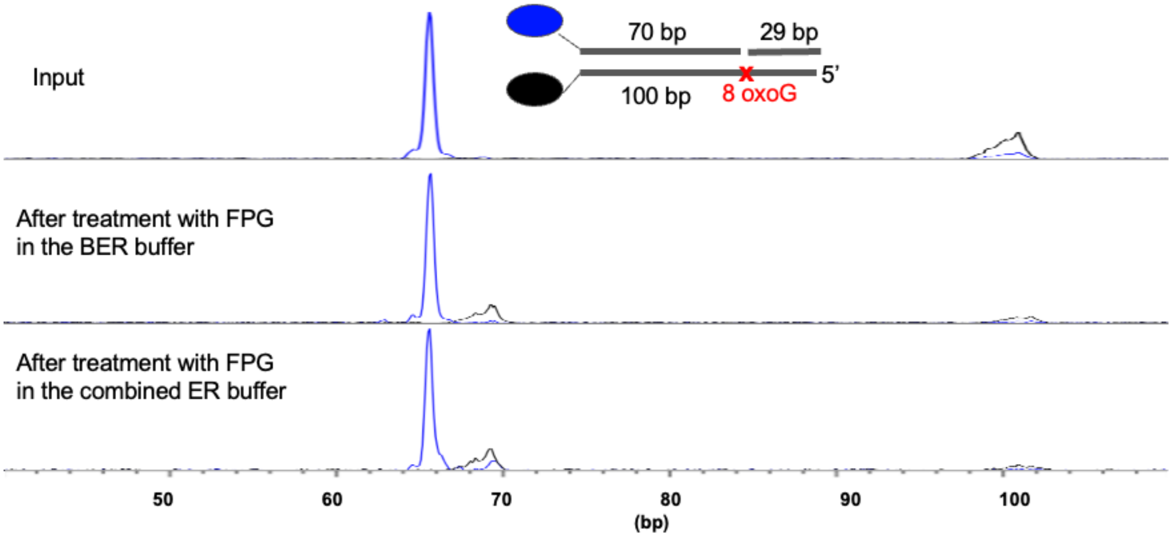


**Figure S11:** Fragment analysis results show that Formamidopyrimidine [fapy]-DNA glycosylase (Fpg) is as active in the integrated ER buffer as in the BER buffer.

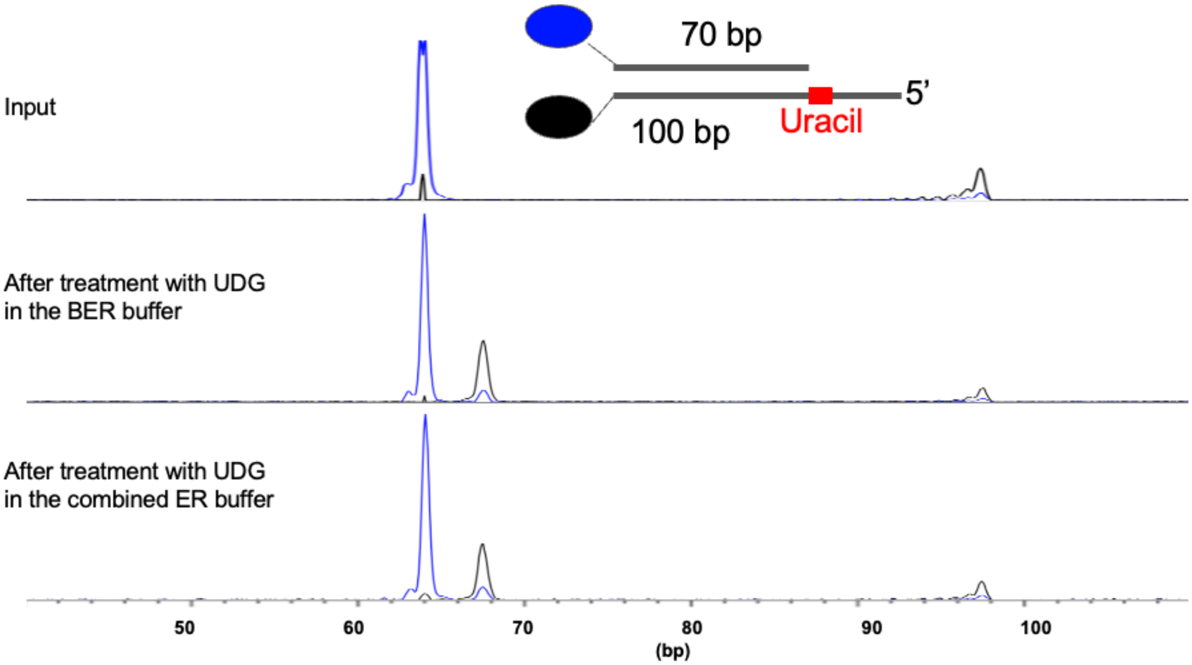


**Figure S12**: Fragment analysis results show that Uracil-DNA glycosylase (UDG) is as active in the BER buffer as in the integrated ER buffer.

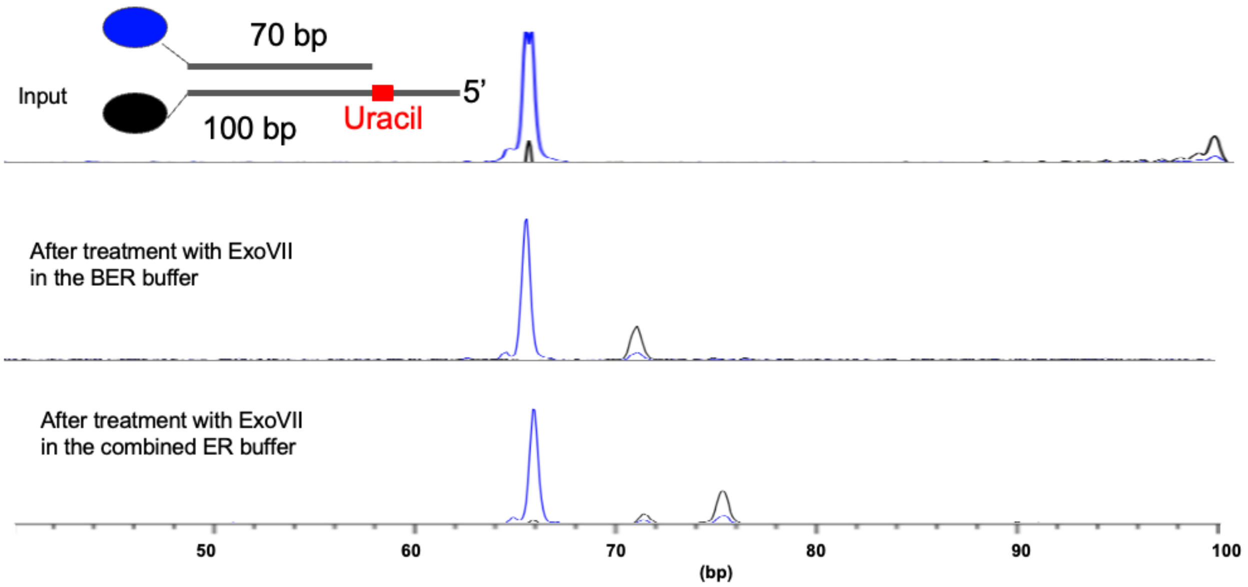


**Figure S13**: Fragment analysis results show that Exonuclease VII (ExoVII) is still active in the integrated ER buffer, though it could leave a slightly longer 5' overhang.

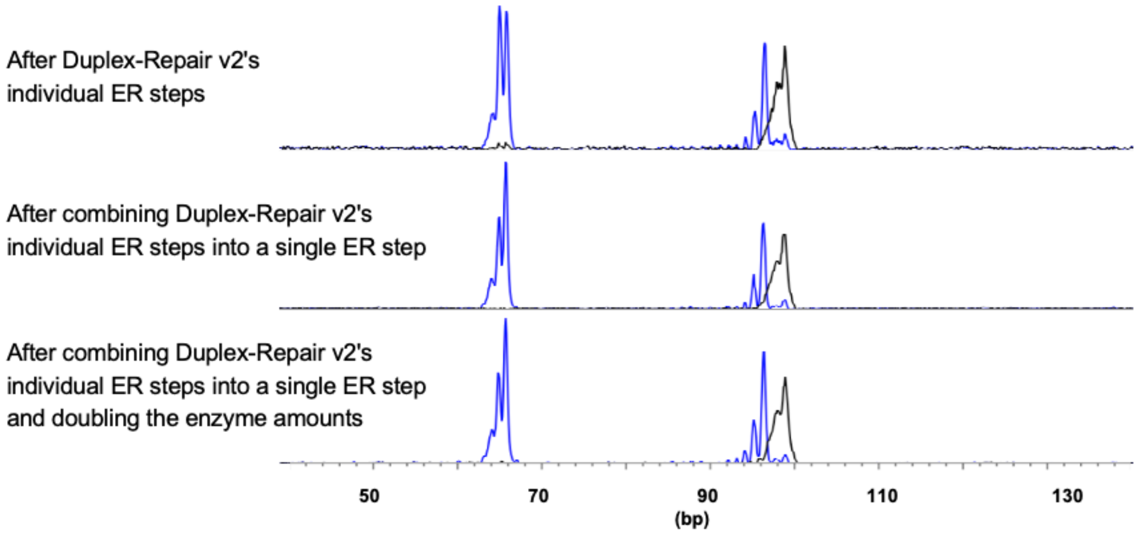


**Figure S14:** Fragment analysis results show combining Duplex-Repair v2's individual ER steps into a single step did not significantly impact the ER efficiency. The same synthetic duplex as in Fig. S10 was used. Of note, the BER was skipped.

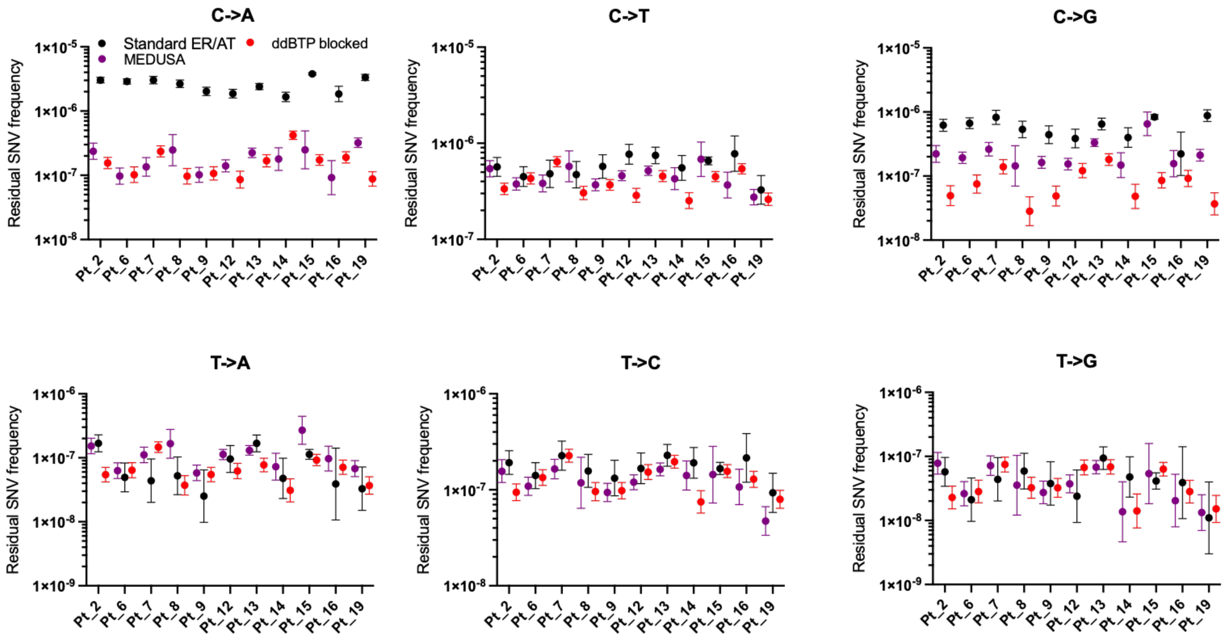


**Figure S15**: Context specific residual SNV frequencies of 11 samples treated with standard ER/AT, MEDUSA, or ddBTP-blocked ER/AT, followed by CODEC whole genome duplex sequencing.

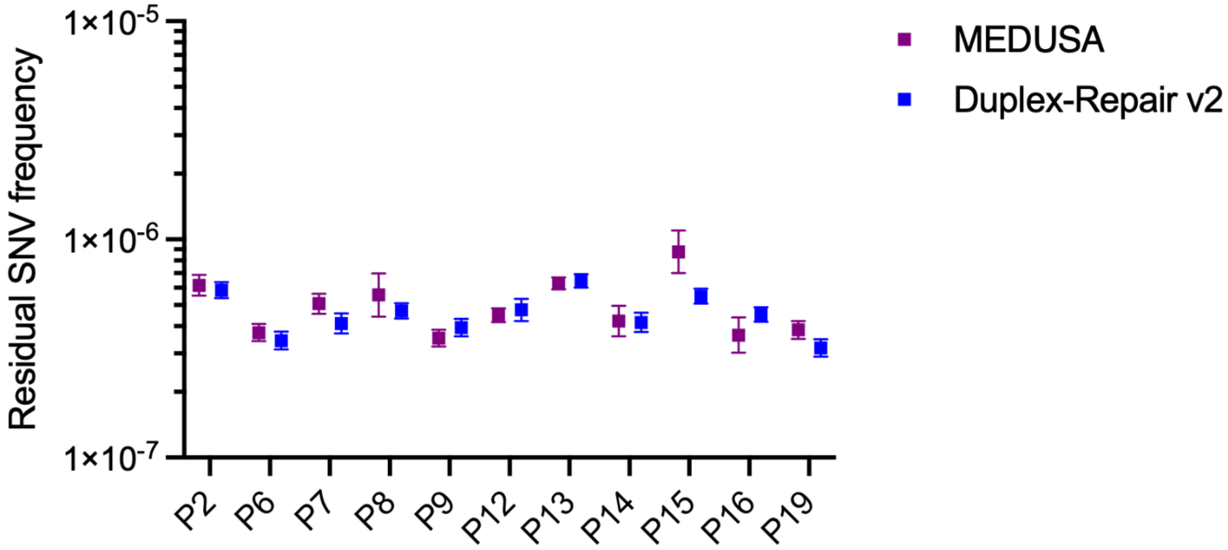


**Figure S16**: Overall residual SNV frequencies of 11 samples treated with MEDUSA or Duplex-Repair v2, followed by CODEC whole genome duplex sequencing.

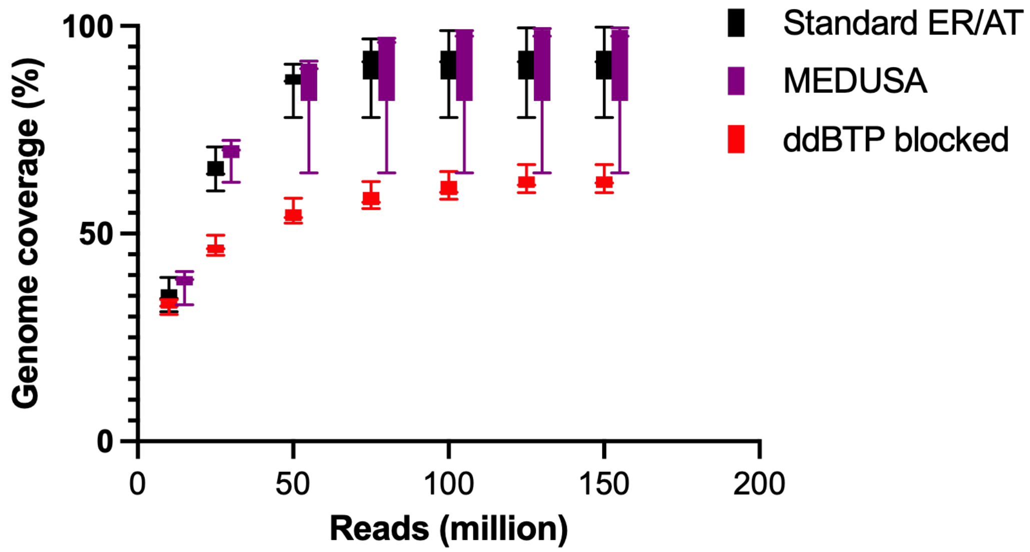


**Figure S17.** CODEC genome coverages of 11 whole blood samples.

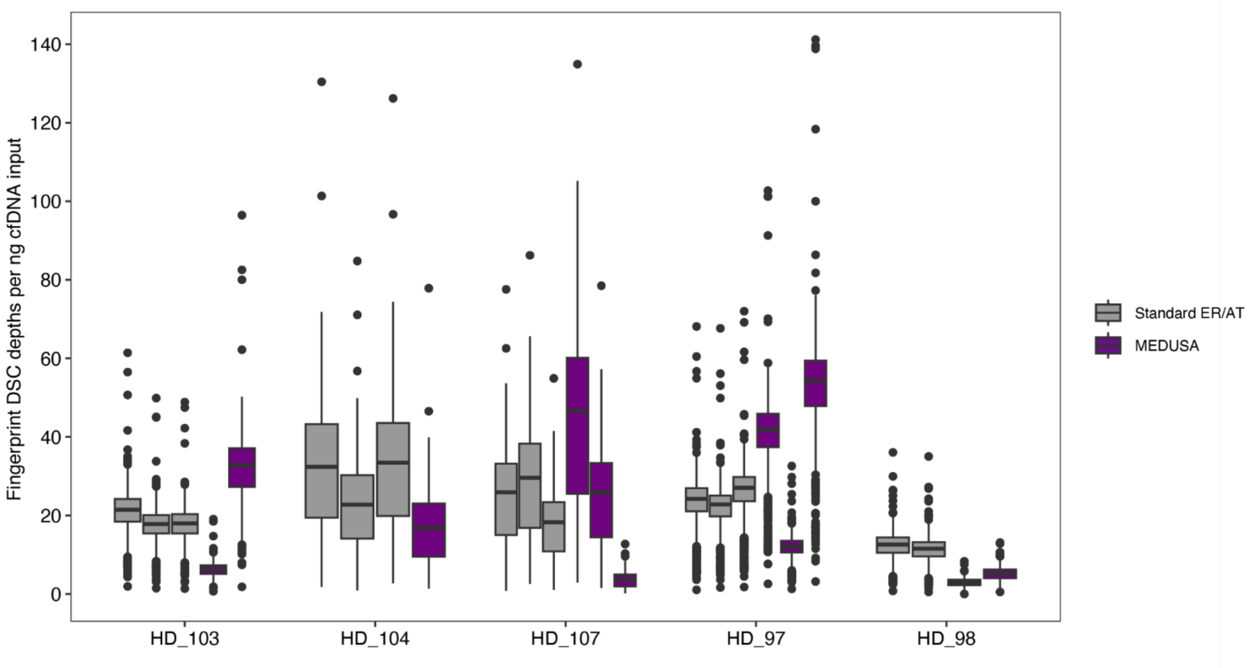


**Figure S18**: Duplex depths recovered of 5 HD cfDNA samples treated with standard ER/AT or MEDUSA, followed by hybrid capture duplex sequencing. We removed replicates that received < 50 million raw reads.

| Oligo ID | Fluorophore end | Fluorophore | Sequence | Length (bp) |
|---|---|---|---|---|
| 1 | 5' | 6-FAM | 6FAM//iSpC3/GCGTCACCAGCCACGCGAGCCGGATGAGGATCCGTGACGCGAAGTCCTGGTACCGCCGCTCGCTTCCGAC | 70 |
| 2 | / | / | GGTTCTCCACCGAGCGACCTAATATTAAT | 29 |
| 3 | 3' | ATTO 550 | ATTAATATTAGGTCGCTCGGTGGAGAACC/**i8oxodG**/GTCGGAAGCGAGCGGCGGTACCAGGACTTCGCGTCACGGATCCTCATCCGGCTCGCGTGGCTGGTGA*C*G*C*/3ATTO550N/(*phosphorothioate bonds) | 100 |
| 4 | 3' | ATTO 550 | ATTAATATTAGGTCGCTCGGTGGAGAACC**U**GTCGGAAGCGAGCGGCGGTACCAGGACTTCGCGTCACGGATCCTCATCCGGCTCGCGTGGCTGGTGA*C*G*C*/3ATTO550N/(*phosphorothioate bonds) | 100 |

**Table S1:** DNA sequences of synthetic oligonucleotides used in this study.
